# Towards the biogeography of butyrate-producing bacteria

**DOI:** 10.1101/2022.10.07.510278

**Authors:** Joel E Brame, Craig Liddicoat, Catherine A Abbott, Robert A Edwards, Jake M Robinson, Nicolas E Gauthier, Martin F Breed

## Abstract

**Aim:** Butyrate-producing bacteria are found in many outdoor ecosystems and host organisms, including humans, and are vital to ecosystem functionality and human health. These bacteria ferment organic matter, producing the short-chain fatty acid butyrate. However, few (if any) studies have examined the macroecological influences on their large-scale biogeographical distribution. Here we aimed to characterise their global biogeography together with key explanatory climatic, geographic, and physicochemical variables.

**Location:** Global, and the Australian continent

**Time period:** 2005-2020

**Major taxa studied:** Butyrate-producing bacteria

**Methods:** We developed new normalised butyrate production capacity (BPC) indices derived from global metagenomic (*n*=13,078) and Australia-wide soil 16S rRNA (*n*=1,331) data, using Geographic Information System (GIS) and modelling techniques to detail their ecological and biogeographical associations.

**Results:** The highest BPC scores were found in anoxic and fermentative environments, including the human and non-human animal gut, and in some plant-soil systems. Within plant-soil systems, roots and rhizospheres had the highest BPC scores. Among soil samples, geographic and climatic variables had the strongest overall influence on BPC scores, with human influence also making key contributions. Higher BPC scores were in soils from seasonally productive sandy rangelands, temperate rural residential areas, and sites with moderate-to-high soil iron concentrations.

**Main conclusions:** Abundances of butyrate-producing bacteria in outdoor soils follow complex ecological patterns influenced by geography, climate, soil chemistry, and hydrological fluctuations. Human population density and soil iron also play substantial roles, and their effects are dependent on a combination of ecological variables. These new biogeographical insights further our understanding of the global ecology patterns of butyrate-producing bacteria, with implications for emerging microbially-focussed ecological and human health policies.

## INTRODUCTION

Butyrate-producing bacteria are both associated with host organisms and are free-living in outdoor ecosystems. They have critical roles in breaking down organic products including fibres (Baxter et al., 2019) and cellulose (Goldfarb et al., 2011). Given suitable organic substrates and anaerobic conditions, these bacteria can produce butyrate, a short-chain fatty acid, as a metabolic by-product of fermentation. In soils, butyrate is associated with the suppression of soil-borne plant pathogens (Poret-Peterson et al., 2019). In humans and non-human animals, gut-associated butyrate provides energy to colonic epithelial cells, maintains gut barrier stability, has anti-inflammatory and immunomodulatory effects, and strengthens the integrity of the blood-brain barrier (Bedford & Gong, 2018; Brame et al., 2021; Knox et al., 2022; Rivière et al., 2016). As such, butyrate production by bacteria is now understood to be a vital part of host-microbe symbioses.

Efforts to quantify the taxonomic and functional abundances of butyrate-producing bacteria through metagenomic analyses have realised multiple challenges. For example, in the SEED functional genome annotation system (Overbeek et al., 2005; https://pubseed.theseed.org/), the subsystem “Acetyl-CoA fermentation to butyrate” (https://pubseed.theseed.org/) includes 20 genes involved in the butyrate production pathway, and individual bacterial genome isolates show varying copies and synteny of the genes for the enzymes (Vital et al., 2014). Quantifying complete functional pathway abundances within samples can therefore be computationally time-consuming and expensive, limiting large-scale sample comparisons. Alternately, molecular methods that directly measure butyrate can provide insights into the functional potential of bacterial communities in samples. However, to our knowledge, few studies have focussed on butyrate in outdoor ecosystems (Brame et al., 2021; P. Liu et al., 2011), and no studies to date have matched soil bacterial short-chain fatty acids and metagenomics on a global biogeographical scale. Therefore, we aimed to develop a novel method to estimate a sample’s potential for butyrate production in a computationally agile way. This workflow could then be scaled up to examine large numbers of samples from a broad spectrum of sources, including outdoor ecosystems.

Soils represent a key reservoir for microbial diversity in ecosystems (Fierer & Jackson, 2006) and provide a focus for us to examine conditions that may support butyrate producers. Soil butyrate producers may be involved in soil-plant-organic matter cycling processes and be ingested and expelled from the digestive tract of soil fauna (e.g., earthworms; Thakuria et al., 2010). At the same time, non-human animal gut-associated bacteria are regularly dispersed into outdoor ecosystems, where they can settle into soils (Blum et al., 2019). Moreover, bacteria in the superficial layer of soils can be dispersed into the air (Polymenakou, 2012; Robinson et al., 2020) and transmitted to hosts (Liddicoat et al., 2020). However, almost no existing studies have quantified outdoor abundances of butyrate-producing bacteria. As such, examining the distribution of butyrate-producing bacteria among the ecological components of their transmission cycle could provide more detailed insight into epidemiological and ecological knowledge of these bacteria, including conditions and mechanisms that support persistence and transmission. Biogeographical knowledge of butyrate-producing bacteria could allow urban landscape architects and managers to understand how certain environmental conditions contribute to the human and non-human exposome. Moreover, it could also allow researchers to develop ways of selecting for potentially health-promoting bacterial assemblages in the environment (Brame et al., 2021).

Here, we examined a broad variety of global sample sources and ecological conditions (e.g., climate, geography, and soil physicochemical characteristics) to identify associations with butyrate production abundances. We developed two new normalised indices to estimate and compare the butyrate production capacity (BPC) of representative samples from both metagenomic (BPC_meta_) and 16S rRNA amplicon (BPC_16S_) data. We combined global metagenomic datasets with continent-wide Australian 16S rRNA amplicon data to provide insight into the global biogeography of butyrate-producing bacteria. Specifically, we aimed to: **(a)** determine the butyrate production capacities across a range of hosts and ecological conditions using our novel BPC_meta_ scores; **(b)** provide a more detailed comparison of BPC_meta_ scores across subcategories within each broader source group; **(c)** conduct an in-depth analysis of Australian soil samples using BPC_16S_ scores to describe ecological associations with soil butyrate production capacity; and **(d)** examine the potential influence of human population density on abundances of soil butyrate producers. By developing novel targeted formulae and collating multiple types of data and analyses, we were able to synthesise a novel and in-depth view into the global biogeography of butyrate-producing bacteria and the interlinkages between their hosts and ecosystems.

## MATERIALS AND METHODS

### Method overview

To compare the bacterial butyrate production capacity of samples across a broad spectrum of hosts and ecosystems, we chose to analyse samples based on two types of bacterial sequencing data: shotgun metagenomic sequences and 16S rRNA amplicons. Each data type required the interrogation of separate databases, using separate statistical tools for analysis and visualisation. We developed novel formulae for each data type to estimate the butyrate production capacity (BPC) for each sample (detailed below). Each formula included a normalisation step to allow BPC scores between samples (within a data type) to be directly compared (**Table 1**).

**Table 1.** Overview of our analyses and data types used.

| Sequenced data type | Shotgun metagenomic sequences | 16S rRNA amplicon sequences |
| --- | --- | --- |
| Formula (see methods for details) | BPC<sub>meta</sub> | BPC<sub>16S</sub> |
| Sample locations | Global (terrestrial and aquatic) | Australia-wide |
| Sample sources | All available | Soil, 1-10 cm depth |
| Database(s) interrogated | Gene data (for *buk* and *atoA* genes) and sample metadata: Integrated Microbial Genomes & Microbiomes | Amplicon data and sample metadata: Australian Microbiome Initiative<br>Ecological data: Atlas of Living Australia, Soil and Landscape Grid of Australia, Socioeconomic Data and Applications Center, and Geoscience Australia<br>Taxonomic data: Genome Taxonomy Database |
| Sample count | Downloaded: $n=22,593$; Cleaned $n=13,078$ | Downloaded: $n=3,023$; Cleaned $n=1,331$ |
| Normalisation step | Single copy gene *pheS* data from IMG/M | Relative abundance |
| Sample BPC score range | 0.02 – 3.39 | 0.01 – 12.59 |
| Statistics and visualisation | After grouping samples into categories, between-group variance and mapping | Principal components analysis, $k$-means clustering, mapping, decision tree modelling |

### Global shotgun metagenomic sample analysis

#### Gene selection and metagenome database interrogation

To characterise the global distribution of butyrate-producing bacteria, we analysed shotgun metagenomic datasets. To begin, the butanoate (butyrate) synthesis pathways were reviewed using Seed viewer subsystems (https://pubseed.theseed.org/) and the KEGG pathway (Kanehisa et al., 2016; https://www.genome.jp/kegg/pathway.html) to determine the genes that code for enzymes that are part of the butyrate production pathway. Based on these pathways, the following two genes were chosen for further analysis: *buk* (butyrate kinase) and *atoA* (acetate-CoA:acetoacetyl-CoA transferase subunit beta). *ACADS/bcd* (butyryl-CoA dehydrogenase) and *ptb* (phosphate butyryltransferase) were analysed but excluded (all gene decisions explained in **Supplementary Table 1** in Supporting Information). The included genes *atoA* and *buk* participate in each of the two terminal pathways. The Integrated Microbial Genomes & Microbiomes (IMG/M) database (Chen et al., 2021) was then searched for each butyrate-related gene to obtain their mean counts among genomes with at least one copy of either gene (**Supplementary Table 2** in Supporting Information). At the time of download, IMG/M utilised Annotation Pipeline v.5.0 protocols (release date October 2018) for functional annotations (Chen et al., 2019).

We next searched global metagenomic databases for *atoA* and *buk* to find metagenomes that suggest the potential presence of butyrate-producing bacteria. Initial gene and translated gene searches of metagenomics data via the ‘Searching SRA’ web facility (https://www.searchsra.org) using bowtie2 and diamond yielded low numbers of samples and/or high E-values. The largest datasets came from searching IMG/M using EC numbers for each butyrate-production enzyme (butyrate kinase = EC 2.7.2.7, acetate-CoA:acetoacetyl-CoA transferase subunit beta = EC 2.8.3.8) as well as three enzymes with single-copy genes used for subsequent normalisation (phenylalanine—rRNA ligase = EC 6.1.1.20; guanylate kinase = EC 2.7.4.8; alanine—tRNA ligase = EC 6.1.1.7). Sample datasets with the genes *atoA* (*n*=21,147) and *buk* (*n*=16,263) were downloaded as our starting point for metagenomics analysis. Among the downloaded datasets, 14,407 datasets had both genes, and many datasets had one gene but not the other (*atoA* only *n*=6,330 and *buk* only *n*=1,856). We then collated these datasets to create an initial set of 22,593 unique metagenomic samples (**Supplementary Table 3** in Supporting Information).

Counts for each butyrate-production gene were normalised by dividing by counts of the single-copy gene *pheS*, which codes for the protein phenylalanine—tRNA ligase alpha subunit and was used as a proxy for total genome count. Counts for two other single-copy genes (*GUK1*: guanylate kinase and *alaS*: alanine—tRNA ligase) were also inspected, but they were not used because *GUK1* searches showed low counts, and *alaS* showed slightly different but proportional counts to *pheS*, which validated the usage of *pheS* to normalise estimates of total genomes in the samples. However, 115 samples did not include *pheS* count data and were subsequently removed from our analysis. Based on examination of data distribution (distribution shown in **Supplementary Figure 1** in Supporting Information), samples with a low *pheS* count <100 (*n*=5,479) and samples with a (*buk*+*atoA)*/*pheS* ratio >30% (*n*=804) were removed from our analysis to minimise bias from samples with a low normalising pheS count. Outlier samples with pheS count >50,000 (*n*=19) were also removed from our analysis. In addition, samples that did not fall within the scope of our research question (*n*=3,098), such as deep subsurface, contaminated (e.g., uranium-contaminated sites), and experimentally altered samples, were excluded from analysis. The remaining 13,078 samples were retained and analysed for our project. The samples originated from a broad range of sources, including soils and sediments, marine samples, human and non-human animal faecal samples, and wastewater samples.

#### BPC scores for shotgun metagenomic samples

To derive the Butyrate Production Capacity (BPC_meta_) score for each sample with metagenomic data, the following formula was developed:

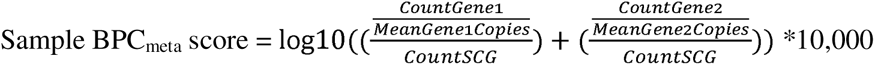

where: SCG = single copy gene (*pheS*)

Gene1 = *buk*, Gene 2 = *atoA*

CountGene1, CountSCG are from global metagenomics sample datasets

MeanGeneXCopies = mean count of copies of gene X among all genomes found from searches of gene X within the IMG/M genome database.

Once BPC_meta_ scores were computed and added to the spreadsheet using Excel formulae, the samples were sorted into six categories: soil and terrestrial sediments, aquatic, human, non-human animal, plant, and agro-industrial (**Supplementary Table 4** in Supporting Information). Samples were then grouped by subcategories for further analysis: human samples were grouped by body compartment; non-human animal samples were grouped by vertebrate/invertebrate and by phylum; plants were grouped by compartment; soil samples were grouped by anthropogenic biome classification (i.e., anthromes, which are “terrestrial biomes based on global patterns of sustained, direct human interaction with ecosystems”; Ellis & Ramankutty, 2008); aquatic samples were grouped by source subcategory; and agro-industrial samples were grouped by source site. To reduce bias within the anthrome “Class” categories, two studies whose samples accounted for >50% of the total class samples were removed from the analysis.

To identify whether our BPC_meta_ formula was estimating butyrate production rather than general anaerobicity, we adapted it to estimate ethanol production, which also requires anaerobic conditions. The butyrate synthesis genes were replaced with the terminal gene for alcohol dehydrogenase (*ADH*, EC 1.1.1.1) to derive an Ethanol Production Capacity (EPC) score. Comparing the resulting EPC scores of the soil metagenomic samples with their BPC_meta_ scores showed a negligible correlation (**Supplementary Figure 2** in Supporting Information). This further validated that our BPC_meta_ scores were specific to butyrate production.

#### Statistics

Statistics were done in R (version 4.0.2; R Core Team, 2020). The Shapiro-Wilk test was used to determine the normality of distribution. In each category the BPC_meta_ scores did not fit a normal distribution, and either the non-parametric Kruskal-Wallis test or the Wilcoxon rank-sum test was used to test for between-group variation. Due to a high *n* in some subgroups, a post hoc Dunn test with Bonferroni correction was used to compare subgroup pairwise differences at α=0.05. The R package ‘ggplot2’ (version 3.3.5; Wickham, 2016) was used for data visualisation. Mapping of samples using the pseudocylindric ‘Robinson’ projection in QGIS (v 3.2.2; QGIS Development Team, 2020) was performed on 2,850 sample metadata coordinates after excluding 360 samples with coordinates with less than two decimal points and 153 samples with no coordinates.

### Australian 16S rRNA amplicon sample analysis

#### BPC scores for 16S rRNA amplicon samples

To assemble 16S rRNA gene abundance data in Australian soil samples, the Australian Microbiome Initiative (Bissett et al., 2016) database was queried for the following parameters: Amplicon = “27F519R”, Kingdom = “bacteria”, Environment = “is soil”, Depth = “between 1 and 10” (cm). We downloaded the abundances of sequences with 100% identity threshold (zero-radius operational taxonomic units or zOTU) and metadata for each resulting sample (*n*=3,023). We used the phyloseq package (McMurdie & Holmes, 2013) for managing and cleaning the 16S rRNA data. We removed all “chloroplast” and “mitochondria” data. We removed low abundance zOTUs that did not occur in at least two samples and had total counts of <20, as these may have arisen from processing errors. In addition, we kept only samples with total number of sequences between 30,000 and 500,000 to remove outliers and samples with low sequence depth. The resulting sample count was *n*=2,795. To normalise the data and reduce bias in comparing samples, we transformed the zOTU abundances into relative abundances.

Because 16S rRNA data can have a relatively poor resolution at the species level (Jovel et al., 2016) as well as many unclassified genera, we focussed our BPC_16S_ derivation on family-level data. Using the Genome Taxonomy Database (GTDB) website interface and the set of putative butyrate-producing species (*n*=118) from Vital et al. (2014), we collated the families that included members in this list of butyrate-producers (*n*=54, **Supplementary Table 5** in Supporting Information). This family list was matched with the Australian Microbiome Initiative taxonomy listings for each downloaded sample. Of the 54 taxonomic families with butyrate-producing bacteria analysed, 31 families had no representative (zOTUs) in any sample. The proportion of butyrate producers in each butyrate-producing family (with *n*=23 such families represented in our dataset) was then used to estimate the abundance of butyrate-producing taxa within each sample and a corresponding BPC_16S_ score, as follows:

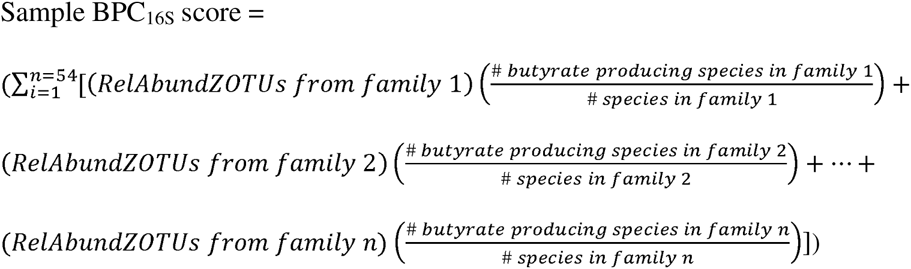

Where:family1= Acetonemaceae, family 2 = Acidaminococcaceae, … (see **Supplementary Table 5** for a list of all 54 families). Count zOTUs in each butyrate-producing family are from Australian Microbiome Initiative datasets. # butyrate-producing species (and total binomial species) in each family were evaluated from the entire GTDB.

#### Modelling ecological associations of BPC_16S_ scores

To provide further biogeographical context to butyrate-producing bacteria in soils, BPC_16S_ scores were associated with geographically-paired ecological metadata. We chose 16S rRNA amplicon-based studies for this analysis because many soil studies in Australia have utilised 16S rRNA data, and the Australian Microbiome Initiative facilitated access to a continental coverage of data collected via consistent sampling and bioinformatic protocols. By selecting a substantial yet manageable spatial scale (i.e., continental Australia), we efficiently examined associations of a larger pool of ecological characteristics with BPC_16S_ scores. We also used ecological metadata from sources focusing solely on Australia (e.g., Atlas of Living Australia), which differs from the metadata sources utilised in our global analyses (e.g., anthropogenic biomes).

To associate ecological data with BPC scores, we first removed any non-continental Australia samples (*n*=342). Covariate data were then collated from a variety of sources and reflected a range of soil-forming factors (i.e., SCORPAN variables: S=soil; C=climate; O=organisms; R=relief; P=parent material; A=age; N=spatial location; McBratney et al., 2003) (see **Supplementary Table 6** in Supporting Information for a list and description of all ecological SCORPAN variables). We identified 49 predictor variables (43 continuous, 6 categorical) as being relevant to our study, for which data sets were downloaded from the following sources: Australian Microbiome Initiative (e.g., organic carbon, clay content %, conductivity; Bissett et al., 2016), Atlas of Living Australia (e.g., annual temperature range, aridity index annual mean; Belbin, 2011; Williams et al., 2010), Soil and Landscape Grid of Australia (e.g., Prescott index, topographic wetness index; O’Brien, 2021), and Geoscience Australia (silica index; Cudahy et al., 2012). We used the best available resolution of source data as supplied to avoid introducing additional noise or bias into our analyses. For example, certain analytical test results were available from sample metadata corresponding to 16S rRNA amplicon data, while other ecological covariate data were extracted from gridded spatial ecological layers at points corresponding to the site locations.

Analysis of the predictor variables showed multiple instances of collinearity (e.g., *r* > 0.80 or Variable Inflation Factor scores >15), and some scatterplots generated showed a curvilinear relationship with BPC_16S_ scores (scatterplots shown in **Supplementary Figure 3** in Supporting Information). Therefore, we used two approaches (detailed below) that were less influenced by collinearity for subsequent analyses: principal components analysis into *k*-means clustering and decision tree modelling via Random Forest (Breiman, 2001). Because many samples did not include soil physicochemical data, removing incomplete cases (*n*=1,122) left 1,331 samples for our analyses (**Supplementary Table 7** in Supporting Information). A further five samples were also removed as they were characterised by outlying values of continuous variables.

We scaled and analysed the 43 continuous predictor variables using principal component analysis (PCA) to reduce the dimensionality of the variables. PC1 and PC2 explained 27.6% and 14.4% of variance, respectively, and were selected for data visualisations (**Supplementary Figure 4** in Supporting Information). We then used *k*-means clustering on scaled original data to assign the samples into clusters. The optimal number of clusters was examined using the “elbow” method, silhouette method, and gap statistic method. While four was considered an optimal number of clusters, we examined both four and five clusters. We found that the additional fifth cluster more distinctly separated land types. Thus, the five-cluster approach was selected for analysis. The resulting cluster data was collated, and BPC_16S_ scores were then matched and returned to the dataset. Median values were calculated for each variable in each cluster, revealing ecological trends distinct to each cluster. Between-cluster significance was examined using Kruskal-Wallis tests. We gave each cluster a generalised description and plotted the sample geospatial coordinates into maps using ‘ggmap’ (Kahle & Wickham, 2013) and Google maps to visualise their geographical distributions.

We then used Random Forest regression modelling (Breiman, 2001) via the R package ‘spatial RF’ (Benito, 2021) to discern variable importance results and obtain partial dependence plots for each variable against BPC_16S_ scores. Only the 43 continuous variables were included in this analysis due to ‘spatial RF’ package limitations. The model fit was estimated using out-of-bag error from the bootstrap. To reduce multicollinearity, highly correlated predictor variables (*r* > 0.8 or VIF > 12) were removed (*n*=9). Tuning the hyperparameters of the model (mtry=24, num.trees=500, min.node size=5) improved its performance (R^2^) by 0.006. Spatial autocorrelation of the residuals was then minimised while fitting the spatial regression model. The resulting Random Forest decision tree model explained 46.5% of the variance in our BPC_16S_ dataset. The variable importance plot was created using random permutations for each predictor variable’s values in out-of-bag data, then calculating the mean decrease in node impurity. Thirty model repetitions were used to create the plot of variable importance. Partial dependence plots were then generated and confirmed the non-linear relationship of most variables with BPC_16S_ scores (**Supplementary Figure 5** in Supporting Information).

## RESULTS

### Global distribution of butyrate-producing bacteria

Metagenomes with genes for butyrate production were found on every continent, in every ocean, and in 89 countries (**Figure 1A**). Overall highest median BPC_meta_ scores were found in human host-associated (2.99, *n*=1,553) and non-human animal host-associated samples (2.91, *n*=771), with the lowest median BPC_meta_ scores in aquatic samples (1.93, *n*=6,017) (**Figure 1B**).

**Figure 1:**
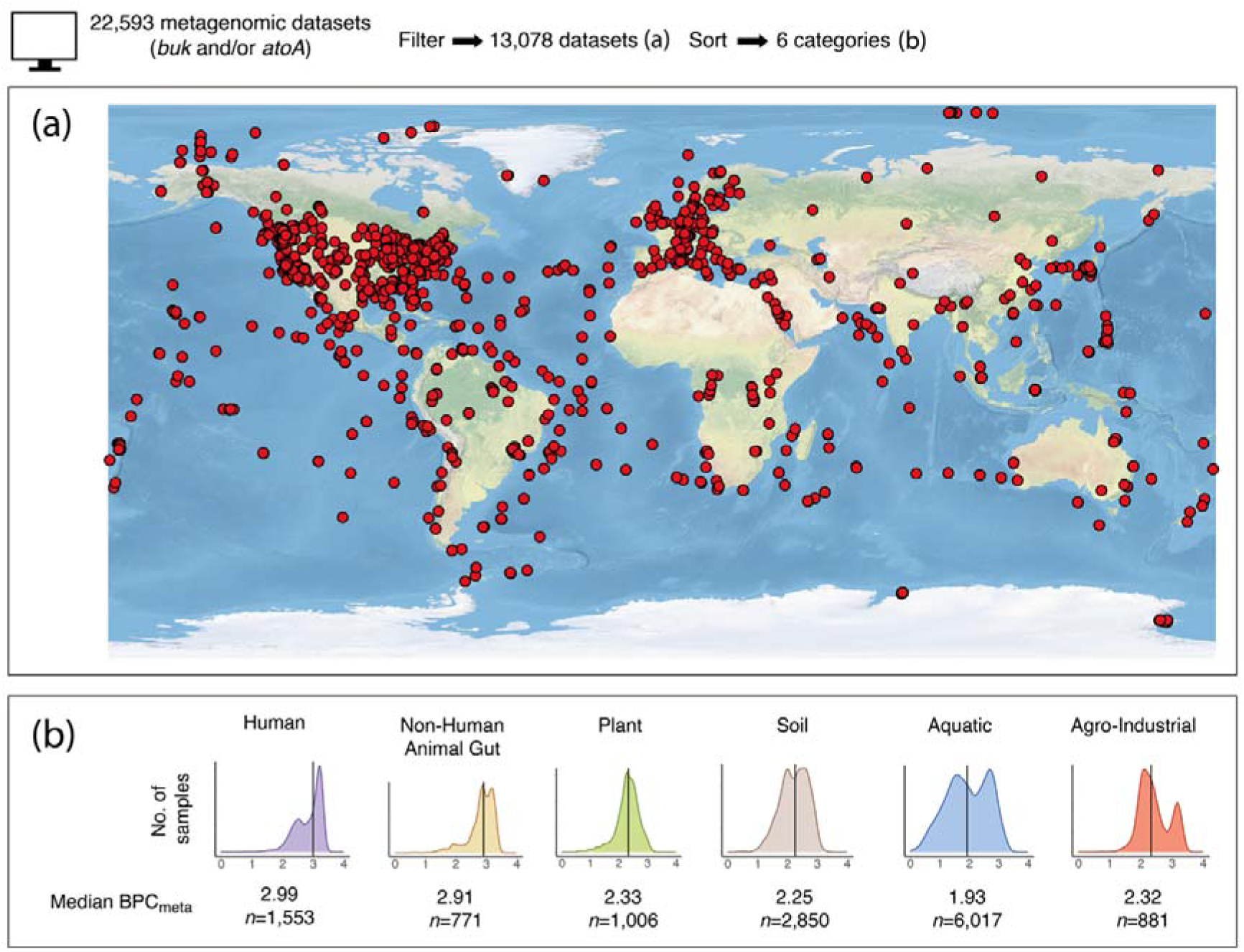
Butyrate-producing bacteria are found on every continent, in every ocean, and in 89 countries. (a) Map showing study locations of samples with *buk* and/or *atoA* genes. (b) Density plots showing frequency distributions of sample BPC_meta_ scores in the six highest-level groupings (x-axis=BPC_meta_). BPC_meta_ score medians rather than means are presented due to non-normal BPC_meta_ score distributions. The range of sample BPC_meta_ scores was from 0.02 to 3.39. Bimodal peaks in five of the six categories may represent divergence between environments supportive and unsupportive of fermentative activity (discussed below). *n* is the number of samples.

### Butyrate production capacity of different ecosystems

#### Human hosts

Human samples were sorted into five body compartments: skin, nasal, oral, genital, and gut. The highest median BPC_meta_ score came from the gut (3.19, *n*=800), with faecal samples expected to be acting as a proxy for the anaerobic gut environment. The lowest median BPC_meta_ score came from the skin (1.86, *n*=17), which is exposed to aerobic conditions. Between-group differences were statistically significant (Kruskal-Wallis test: H=1136, 4 d.f., *P*=<0.001; **Figure 2A**).

**Figure 2:**
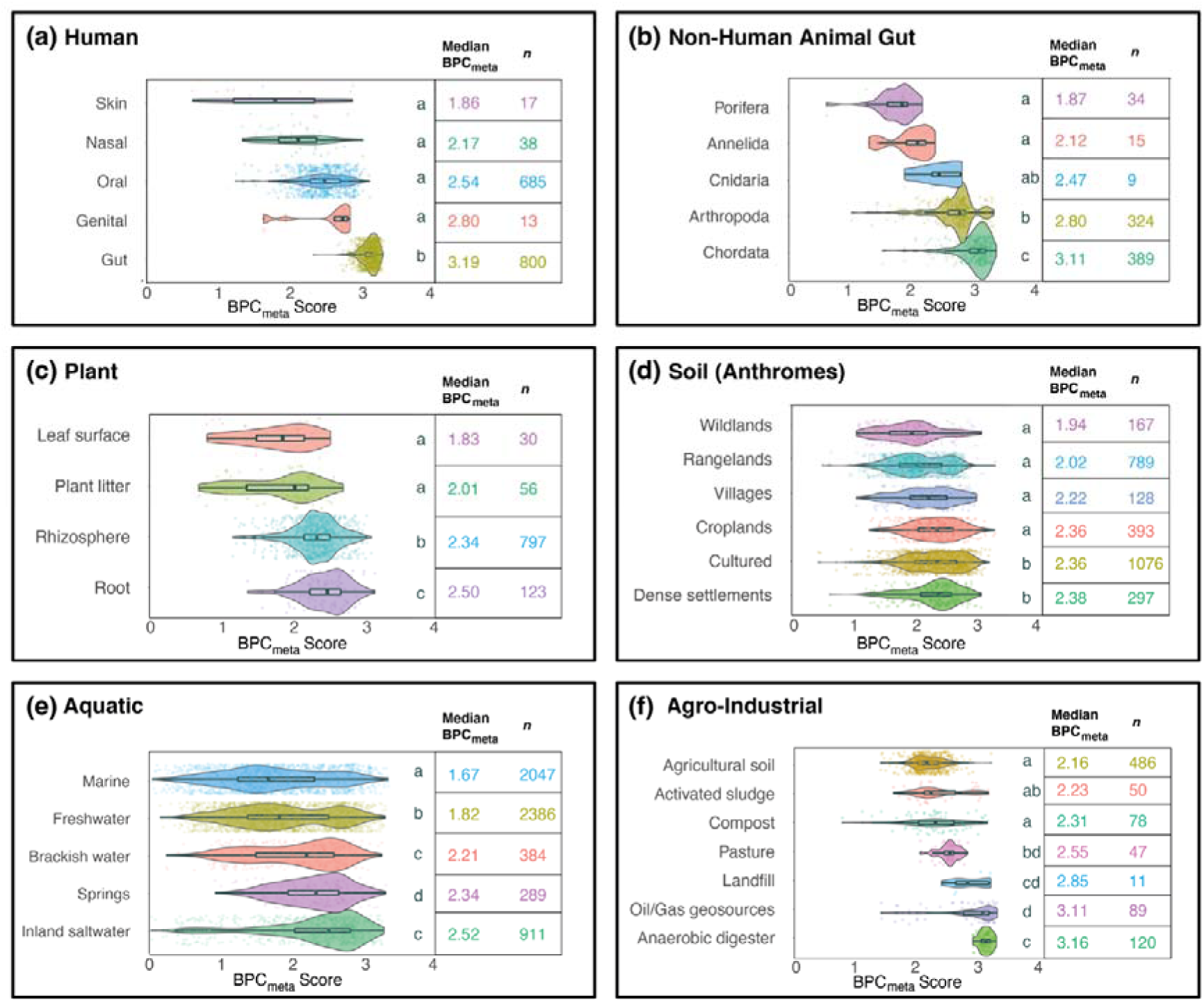
BPC_meta_ scores vary between host communities and ecological sources. BPC_meta_ score density plots by group subcategories. (a) BPC_meta_ scores of humans, sorted by body compartment. (b) BPC_meta_ scores of non-human animal-associated microbial communities, sorted by class. Note that Porifera do not possess a gut. (c) BPC_meta_ scores of plant-associated samples, grouped into compartments. (d) BPC_meta_ scores of soil samples, grouped into anthropogenic biomes (anthromes) levels that represent human influence on land use. The level “Cultured” includes woodlands and inhabited drylands. (e) BPC_meta_ scores of aquatic ecosystem samples, grouped into source site categories. (f) BPC_meta_ scores of agricultural and industrial samples, grouped by source site. Activated sludge and anaerobic digesters are common components of wastewater treatment plants. In each of (a)-(f), Kruskal-Wallis tests show that between-group differences were significant at P<0.001. Medians sharing a letter are not significantly different by the adjusted Dunn test at the 5% significance level. Boxes show the interquartile range.

#### Non-human animal hosts

Non-human animal host-associated samples included in our analysis (*n*=771) were either direct or proxy (e.g., faecal) measures of animal gut microbiota (*n*=448) or were non-gut but host-associated samples (e.g., attine ant fungus gardens, gutless marine worms, *n*=323). We first compared non-human animal groupings by vertebrates (median BPC_meta_ score=3.11, *n*=389) and invertebrates (median BPC_meta_ score=2.76, *n*=382) (between-group differences were statistically significant: Wilcoxon rank sum test: W=22,592, *P*<0.001). We then compared non-human animal samples by taxonomic phylum (between-group differences were statistically significant: Kruskal-Wallis test: H=331, 4 d.f., *P*<0.001; **Figure 2B**), where Chordata had the highest median BPC_meta_ score (3.11, *n*=389) and Porifera (sponges), which lack a gut, had the lowest BPC_meta_ scores (1.87, *n*=34). A further comparison showed that the human gut median BPC score (3.19, *n*=800) was similar to the primate gut (3.12, *n*=26).

#### Plant hosts

Our dataset included 1,006 plant-associated samples. These were subcategorised into four groups by plant compartment: leaf surface, plant litter, rhizosphere, and root. Root samples had the highest median BPC_meta_ score (2.50, *n*=123). Leaf surface samples had the lowest median BPC_meta_ score (1.76, *n*=30; between-group differences were statistically significant: Kruskal-Wallis test: H=105, 3 d.f., *P*<0.001; **Figure 2C**).

#### Soil ecosystems

Soil samples (*n*=2,850) were sorted using the anthropogenic biome (anthrome) categories (Ellis et al., 2021; Gauthier et al., 2021), representing varying densities of human population and land use (anthrome classes and world map shown in **Supplementary Figure 6** in Supporting Information). At the “Level” category of anthromes, the highest median BPC_meta_ scores came from both “Dense settlements” (includes classes “urban” and “mixed settlements”; median BPC_meta_ score=2.38, *n*=297) and “Cultured” (includes “woodlands” classes and the “inhabited drylands” class; median BPC_meta_ score=2.36, *n*=1076). The lowest median BPC_meta_ score (1.94, *n*=167) came from the anthrome level “Wildlands”, which has the lowest human influence (between-group differences were statistically significant: Kruskal-Wallis test: H=186, 5 d.f., *P*<0.001; **Figure 2D**).

#### Aquatic ecosystems

Aquatic samples (*n*=6,017) were sub-grouped into five categories: marine, freshwater, brackish water and estuary, springs, and inland saltwater. The highest median BPC_meta_ score (2.52, *n*=911) was found in inland saltwater samples, and marine samples had the lowest median BPC_meta_ score (1.67, *n*=2047) (between-group differences were statistically significant: Kruskal-Wallis test: H=530, 4 d.f., *P*<0.001; **Figure 2E**).

#### Agricultural and industrial samples

Agricultural and industrial (“agro-industrial”) samples (*n*=881) were from a wide variety of sources and materials. We grouped them into seven source types, which include two sample types from wastewater treatment plants (i.e., activated sludge from aeration tanks and anaerobic digesters). The highest median BPC_meta_ scores (3.16, *n*=120) were from anaerobic digester samples. The lowest median BPC_meta_ scores were from the agricultural soils (2.16, *n*=486) and activated sludge (2.23, *n*=50) (between-group differences were statistically significant: Kruskal-Wallis test: H=431, 6 d.f., *P*<0.001; **Figure 2F**). Activated sludge is a bacteria-rich product formed in aeration tanks with aerobic conditions.

### Ecological characteristics associate with soil BPC scores

When clustering the Australian 16S rRNA soil samples on ecological metadata, each cluster aligned with distinct representative land types, which were given the following descriptive titles: ‘arid inland clay plains’ (cluster 1), ‘seasonally productive sandy soils’ (cluster 2), ‘sandy inland deserts’ (cluster 3), ‘temperate urban hinterland’ (cluster 4), and ‘wet, cold, acidic, vegetated montane’ (cluster 5). Mapping the geographical locations of the sample sites showed consistency with the clustered land type descriptions (**Figure 3A**). Median BPC_16S_ scores varied significantly between the clusters (Kruskal-Wallis test: H=164, 4 d.f., *P*<0.001; **Figure 3B**). The highest median BPC_16S_ scores came from the ‘seasonally productive sandy soils’ (0.93, *n*=383) and ‘temperate urban hinterland’ cluster (0.86, *n*=440). The lowest median BPC_16S_ score came from the ‘wet, cold, acidic, vegetated montane’ cluster (0.49, *n*=208). Principal components analysis of all continuous ecological predictor variables showed two distinct patterns of axial distribution, ecological wetness and greenness and soil fertility, and dimensions 1 and 2 explained 27.6% and 14.4% of the variation in the data, respectively (**Figure 3C**).

**Figure 3:**
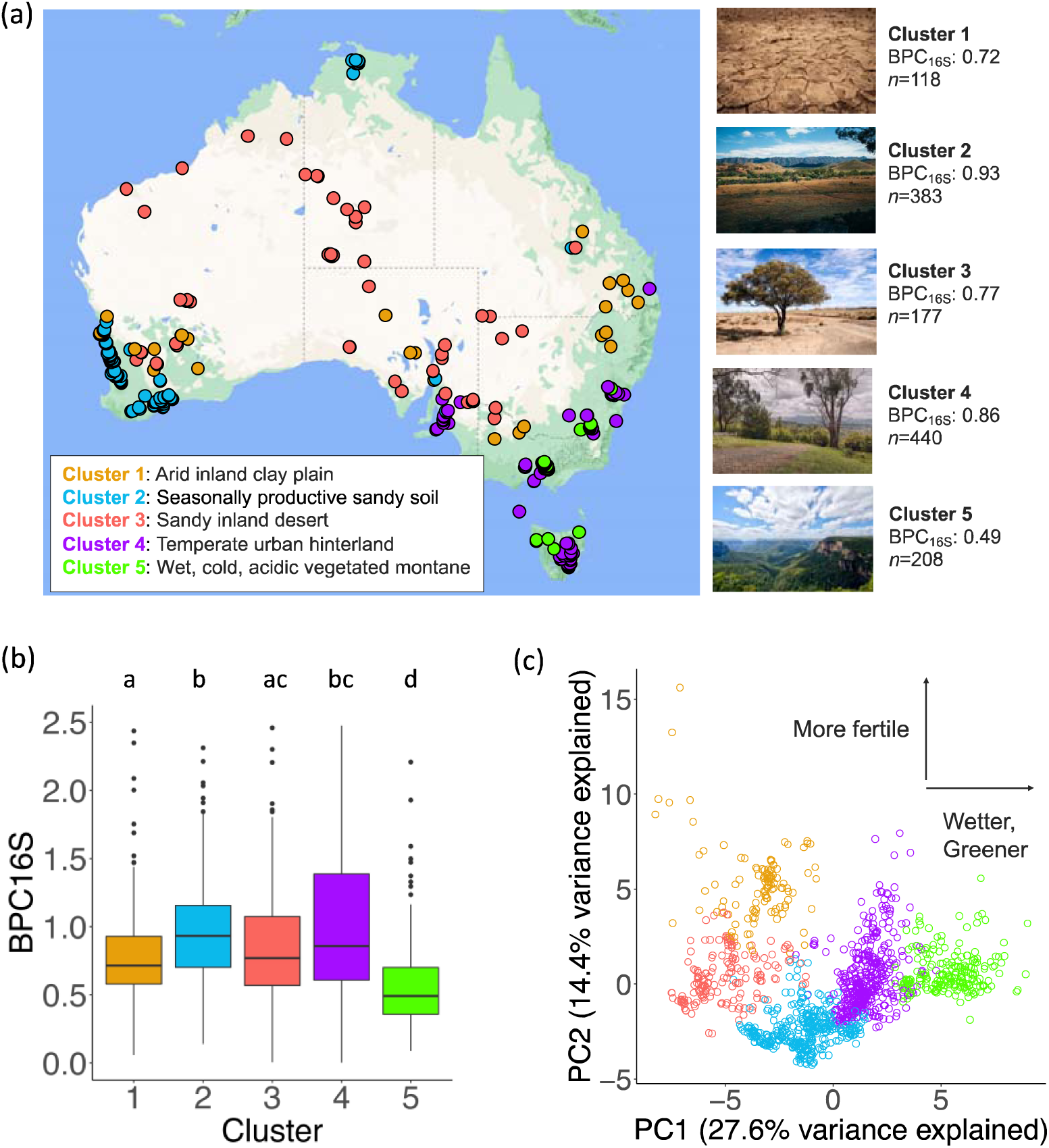
Clustering of ecological data shows five distinct land types. (a) Map of Australian soil samples clustered on 43 continuous ecological variables, five cluster distribution, mapped using R package ggmap and Google maps. Photographs were downloaded from Unsplash.com under CC0 license. (b) Boxplots of median BPC_16S_ scores across each of the five clusters. Medians sharing a letter are not significantly different by the adjusted Dunn test at the 5% significance level. Boxes show the interquartile range. (c) The first two principal components coloured by k-means clusters. The x-axis can be broadly interpreted as ecological wetness and greenness and associated variables (e.g., vegetation cover). The y-axis can be broadly interpreted as soil fertility and the presence of cations.

The cluster with the highest median BPC_16S_ score was the ‘seasonally productive sandy soil’, which has low clay content, low levels of cations, high precipitation seasonality and mean temperature, and generally moderate topographic relief and Prescott index (a measure of soil water balance based on average precipitation and potential evaporation). The cluster with the second-highest median BPC_16S_ score was ‘temperate urban hinterland’, which is generally moderate in elevation, annual rainfall, topographic relief, clay, and soil fertility and has high levels of zinc and manganese. The cluster with the lowest median BPC_16S_ score, ‘wet, cold, acidic, vegetated montane’, had high elevation and topographic relief, colder mean annual temperature, high annual rainfall and aridity index, consistent rainfall levels throughout the year, high soil organic carbon content, and high soil iron and aluminium content (where soil aluminium content and pH are inversely correlated). The two additional clusters, ‘arid inland clay plains’ and ‘sandy inland desert’, also had distinct characteristics (**Supplementary Table 8** in Supporting Information). A separate analysis of categorical variables also showed the highest BPC_16S_ scores among specific land cover types (“built-up” and “plantation”), land use types (“grazing of native pastures” and “rural residential”), and anthromes (“mixed settlements” and “remote rangelands”) (**Table 2** and **Supplementary Table 9** in Supporting Information).

**Table 2.** Categorical variable subcategories with the highest BPC_16S_ scores*.

| Category | Highest median BPC<sub>16S</sub> | Second-highest median BPC<sub>16S</sub> |
| --- | --- | --- |
| **Land cover** | Built-up (1.39, *n*=53) | Plantation (softwood/mixed) (1.13, *n*=11) |
| **Land use** | Grazing of native pastures (1.39, *n*=439) | Rural residential (1.28, *n*=26) |
| **Anthrome** | Mixed settlements (1.07, *n*=28) | Remote rangelands (1.00, *n*=1,103) |
| **Vegetation type** | Eucalyptus open woodlands (1.48, *n*=147) | Hummock grasslands (1.34, *n*=77) |
| **Major vegetation subgroup** | Mitchell grass (*Astrebla*) tussock grasslands (1.90, *n*=38) | Eucalyptus open woodlands with a grassy understorey (1.80, *n*=28) |

<sup>\*</sup> Only subcategories with *n*≥10 are reported here.

The variable importance plot from Random Forest decision tree analysis showed that geography most closely associates with the BPC_16S_ scores in soils (**Figure 4A**). The eight top predictor variables were longitude, elevation, iron, topographic relief, latitude, precipitation seasonality, population density, and the Prescott index. Specifically, low-to-moderate elevation, higher soil iron, and moderate-to-high topographic relief showed a close relationship with higher BPC_16S_ scores (**Figure 4B**). Partial dependence plots for each variable are shown in **Supplementary Figure 5**.

**Figure 4:**
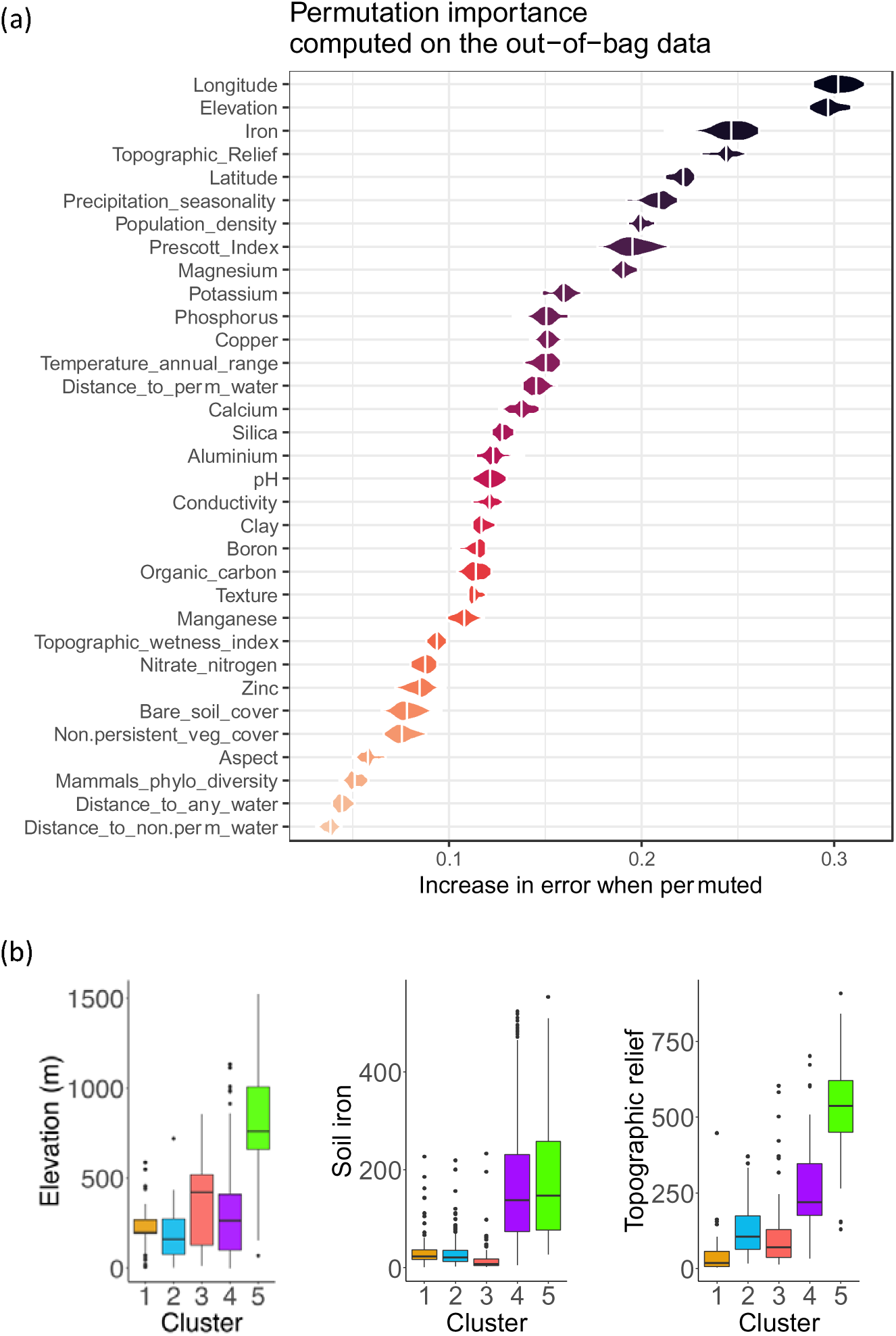
Variable importance results from Random Forest decision tree modelling. (a) Random Forest variable importance results from 43 continuous ecological predictor variables. The model was fitted using out-of-bag errors from the bootstrap. The variable importance was determined using random permutations of predictor variables and the mean decrease in node impurity. (b) Boxplots for selected top variables from (a) across each of the five clusters. In each of elevation, soil iron, and topographic relief, Kruskal-Wallis tests show that between-group differences were significant at *P*<0.001. Boxes show the interquartile range.

## DISCUSSION

We report ecological and biogeographical patterns of butyrate-producing bacteria via our novel BPC formulae that use both metagenomic and 16S rRNA amplicon bacterial data. Our BPC_meta_ data showed that samples from anaerobic and fermentative conditions, such as the animal gut compartment and soil rhizosphere, had increased genomic potential for butyrate production. Our BPC_16S_ results showed that geographical and climatic variables, soil iron, and human population density were key ecological covariates of soil butyrate production capacity. Soils from sandy, sparsely-populated rangelands with high precipitation seasonality, as well as temperate peri-urban sites with greater soil fertility, had higher BPC_16S_ scores. We discuss the novel and distinct ecological and biogeographical patterns of butyrate production observed below.

### Anaerobic conditions associated with higher butyrate-production gene abundances

We show that samples from the chordate gut and anaerobic digesters had the highest BPC_meta_ scores. These results are consistent with the knowledge that butyrate production occurs in anaerobic compartments, such as the human gut and anaerobic digesters (Conrad, 2020; C. Liu et al., 2016). Expectedly, air-exposed skin surface, animals without a digestive system (e.g., sponges), and activated sludge from aeration tanks had among the lowest BPC_meta_ scores for their respective categories. However, our results also show that this anaerobic requirement is maintained within plant-soil systems that are more susceptible to complex ecological dynamics. Underground compartments, such as roots and rhizospheres, had significantly higher median BPC_meta_ scores than air-exposed compartments such as leaf surfaces. Uteau et al. (2015) demonstrated within plant rhizospheres a spatial oxygen gradient based on the presence of oxygen-filled pores. They also found that episodic water-saturated conditions within the rhizosphere decreased the oxygen partial pressures substantially, which may help explain our finding that topographic relief, precipitation seasonality, and evapotranspiration rates play important roles in the butyrate production capacity within soils.

### Influence of human population density on soil BPC_16S_

We report an association between the population density of humans and soil BPC_16S_ scores. Soils from the ‘built-up’ land cover, ‘mixed settlements’ anthrome, and ‘temperate urban hinterlands’ cluster showed among the highest BPC_16S_ scores. In contrast, soils from the ‘remote woodlands’ and ‘wild woodlands’ anthromes had among the lowest median BPC_16S_scores. The anthropogenic influence on terrestrial ecosystems is well-known (Ellis et al., 2021), and our findings show this influence extends to microbial abundances, particularly butyrate-producing bacteria.

We show that ‘temperate urban hinterlands’ are a substantial reservoir of butyrate-producing bacteria. Geographical and climatic data show that these Australian sample sites are near major cities and are moderate in their elevation, mean temperature, topographic relief, and annual rainfall. Thus, the rainfall tends to run off rather than accumulate, which creates the potential for fluctuating hydrology. Additionally, the increased BPC_16S_ scores from these peri-urban locations may be influenced by inadvertent dissemination and artificial soil ‘contamination’ with gut-associated bacteria from people and their pets (Blum et al., 2019), possibly contributing into a transmission cycle of butyrate producers. On the other hand, Australia’s major cities and hinterlands are typically coastal, often with river-floodplain systems and areas of fertile soils that were attractive to European settlers. Thus, our BPC_16S_ results suggest an association between high human population density and high BPC scores. However, the direction of influence remains a compelling question that may be of future research interest.

### Hot, seasonally productive sandy soils

Our cluster ‘seasonally productive sandy soils’ had the highest BPC scores. These soils associated with high precipitation seasonality, which was the climatic variable with the strongest importance toward BPC scores. Hydrological fluctuations in soils have been shown to induce changes in both bacterial community structure and exometabolites (extracellular metabolites). For example, the wetting process in dry soil biocrusts can induce a shift from a cyanobacteria-dominated to a Firmicutes-dominated community within 49.5 hours (Swenson et al., 2017). RoyChowdhury et al. (2022) also reported that soil abundances of Firmicutes increased in a dry-to-wet transition, even more so than in fixed saturated conditions. Because most butyrate producers are within the phylum Firmicutes, hydrological fluctuations may play a key role in butyrate production.

The ‘seasonally productive sandy soils’ cluster also had a high mean temperature, low soil fertility, and low population density, which differs from the above patterns of higher BPC scores around temperate residential areas. Furthermore, these soils had high non-persistent vegetation cover (**Supplementary Table 8**). Seasonal rainfall could lead to flushes of green growth, followed by a suitable climate to enable microbial breakdown. Sandy soils would indicate less protection of organic matter associated with clay minerals, which could enhance microbial decomposition of existing organic matter (Johnston et al., 2009). This suggests a high turnover system, and indeed this cluster contains 67% of our samples characterised by land use as ‘cropping lands’ (**Supplementary Table 7**).

### Soil iron associated with BPC

We observed that iron had the strongest association with BPC scores. Our partial dependency plot shows moderate soil iron levels associated with decreased BPC scores, but BPC scores then increased with higher iron levels. This result is consistent with findings from Dostal et al. (2015), where high iron levels enhanced butyrate production in *Roseburia intestinalis* cultures. Additionally, the authors’ *in vitro* child gut fermentation model showed that a strong iron deficiency significantly decreased butyrate production. It is worth noting that the butyrate production pathway requires several ferredoxins and ferredoxin-like proteins, which are proteins structured with iron, during the reduction of crotonyl-CoA into butyryl-CoA (Chowdhury et al., 2015). On the other hand, our results also show that ‘wet, cold, acidic montane’, the cluster with the lowest median BPC_16S_ score, also had a high concentration of iron. Lee et al. (2001) found that butyrate production from a sucrose solution began to decrease as iron concentration increased above 20 mg/l. Iron availability is also pH dependent, increasing in acidic conditions. Thus, the relationship between iron concentrations and butyrate production is not linear and may rely on a suite of conditions, possibly including the presence of wetting/drying cycles, that work in tandem to create the potential for butyrate production.

### Future research directions and limitations

During the development of our methods, several limitations of our study became apparent. Analyses of shotgun metagenomic sequences and bacterial 16S rRNA amplicons rely on reference databases that are continually being developed but are incomplete. Missed or incomplete sequence identification could affect the reliability of our formulae. Likewise, taxonomy databases are regularly updated, but their updates are not uniform across databases. We used the Genome Taxonomy Database to classify our list of butyrate-producing bacteria, but it showed occasional discrepancies with the classification system applied to the Australian soil 16S rRNA data. Thus, utilising multiple taxonomic classification systems likely means that some butyrate producers were not identified in our data. This could affect the reliability of our results and a future sensitivity analysis is warranted.

To maximise the precision of our butyrate producer database, we chose to use species-level classifications via GTDB representative species. However, this may have inadvertently created inconsistent data from species with multiple strains (sometimes hundreds of strains are present within a species), among which some may be butyrate producers and others not. Analysis at the strain level could provide a higher resolution of data, which should be a future research priority. In addition, our data were genomic only and did not include direct measurements of butyrate concentrations; this would require different data types (e.g., metabolomic data) and pipelines. Future studies that examine butyrate concentrations in relation to butyrate-producing taxonomic and functional gene abundances could more precisely reveal conditions that promote active butyrate production beyond our estimations. Furthermore, because our 16S rRNA data came only from Australia, our modelling may not be generalisable to global conditions that exceed the ranges of our ecological covariate data. For example, the height of mountains on the Australian mainland does not exceed 2,228 m; thus, our mountain cluster modelling may not fit other countries with higher mountains.

In our examination of soil shotgun metagenomic data using anthromes, the sampling methods were not disclosed and might not have been uniform. Metadata that includes sampling depth or indicates whether the soil is horizontally layered or horizontally homogenised would be necessary to draw more comprehensive conclusions. There is also a possibility that urban soils may also be influenced by either the use of compost, which may still bear bacterial spores or DNA (Subirats et al., 2022), or direct contamination by animal faeces. Thus, we have presented the soil BPC_meta_ results but have refrained from drawing conclusions from them; future studies should include details of the sampling methods, which would reduce such biases.

Finally, our data depends on the capacity of laboratory DNA extraction methods to open endospores. Sampling methods that expose the samples to air may inadvertently cause the sporulation of bacteria. Such methods may subsequently reduce the quantities of DNA extracted from spore-formers, several of which could be butyrate-producing bacteria (Browne et al., 2016). Consistency across sampling and DNA extraction methods in future studies could help improve butyrate-producing bacterial abundance data reliability.

## Conclusions

Butyrate-producing bacteria provide critical ecosystem services for hosts and environments, including humans, and soils. Our study focussed on the distribution and ecological associations of these bacteria, building new knowledge of their roles in human-plant-soil ecosystem dynamics. We present new evidence that many outdoor ecosystems influenced by human-associated processes may represent key reservoirs of butyrate producers. Because nearly 60% of the world’s population now lives in urban areas (Güneralp et al., 2020), understanding the influence of dense populations of humans on outdoor urban microbiomes is essential to biodiversity research, informing urban landscape design, and the study of biodiversity-human health linkages (DelgadolJBaquerizo et al., 2018; Kondo et al., 2018; Watkins et al., 2020). Our study helps advance these research areas. Future assessment of butyrate-production capacity across fine spatial scales (i.e., below global and continental, as used here) will help provide greater detail for city infrastructure planning and further microbiome-based public health and ecology research.

## Data availability

The datasets generated during and/or analysed in the current study are available in Supplementary information, and all datasets and custom R code are available on figshare at https://figshare.com/s/3ae2f1327a11f91793d8.

