## Supporting Information for "Towards the biogeography of butyrate-producing bacteria"

Running Title: Biogeography of butyrate-producing bacteria

**Keywords**: butyrate, ecosystem services, gut microbiome, metagenomics, microbial ecology, soil microbiota

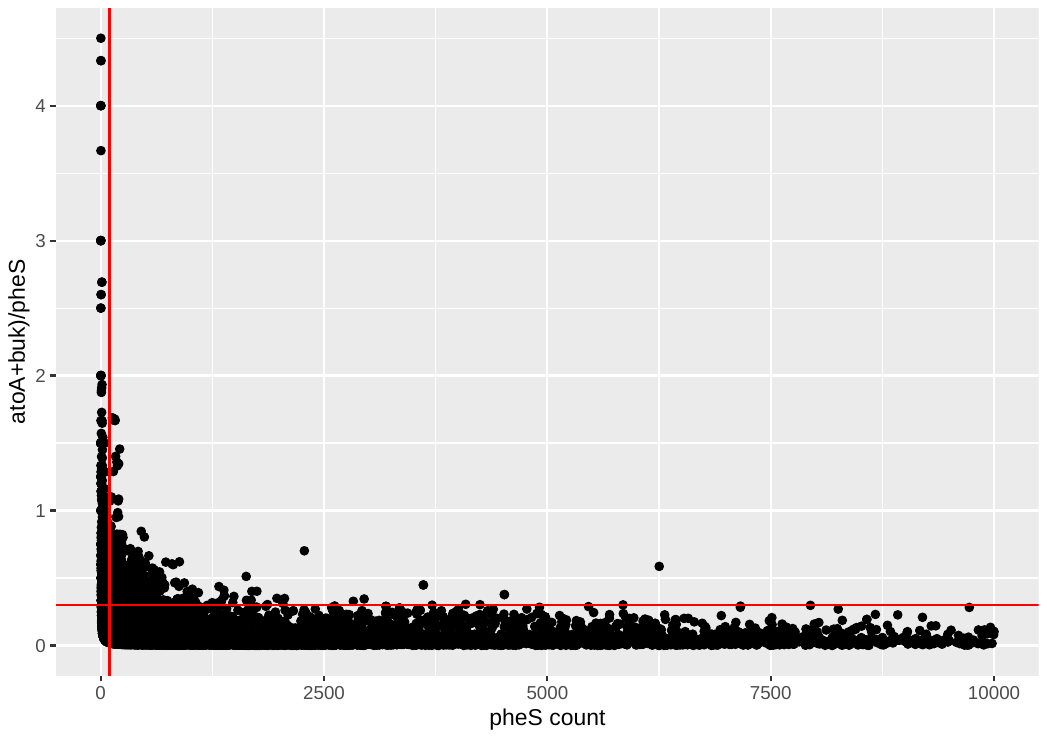

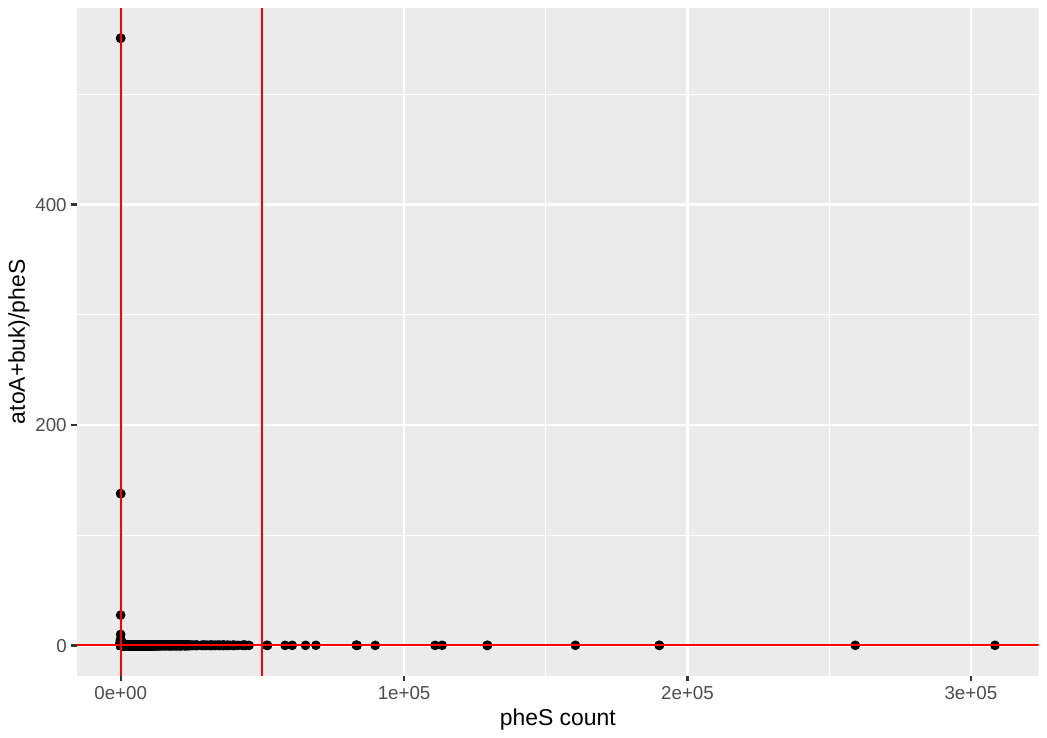

(b)

(a)

**Supplementary Figure 1. Scatterplots showing thresholds of pheS counts and (atoA+buk)/pheS ratio.**

Scatterplots of pheS count samples against (atoA+buk)/pheS ratios. (a) The horizontal red line shows the threshold (y-intercept=0.3) of the (atoA+buk)/pheS ratio, above which samples were removed from analysis. The vertical red lines show the thresholds (x-intercepts 100 and 50,000) of the pheS counts, outside of which samples were removed from analysis. (b) Data from (a) shown with x-axis upper limit=10,000 and y-axis upper limit=5.0 set for a clearer visualisation of thresholds.

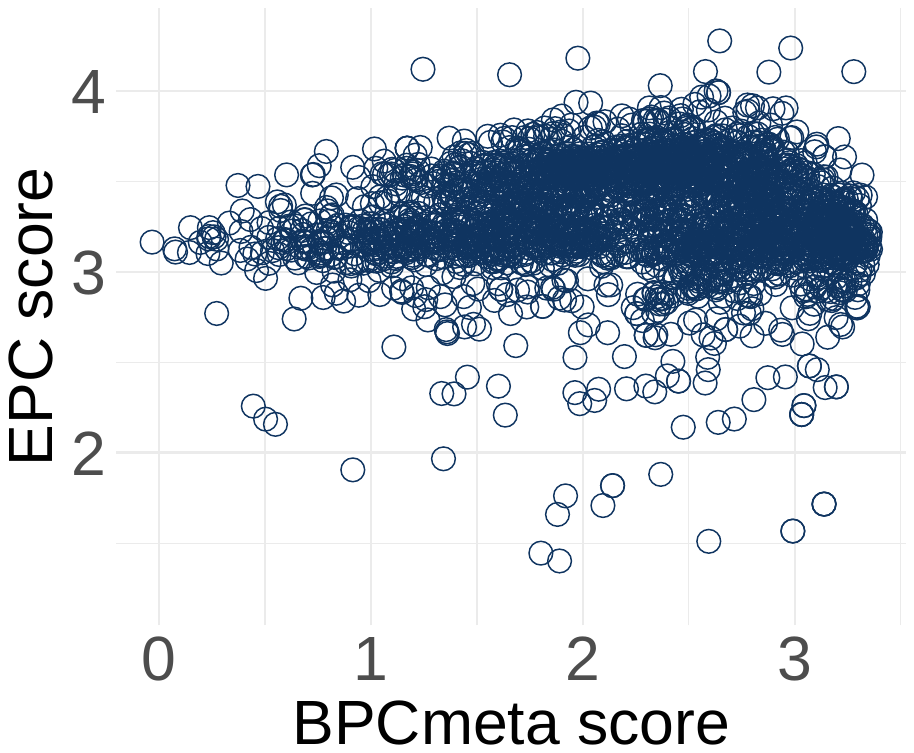

**Supplementary Figure 2. Butyrate production capacity scores show specificity toward butyrate production rather than general anaerobicity.**

Scatterplot of two anaerobic condition production capacities of soil bacteria, ethanol production capacity (EPC) and butyrate production capacity (BPC_meta_) scores, among metagenomic soil samples (*r*^2^=5.6e-05, *p*=0.70).

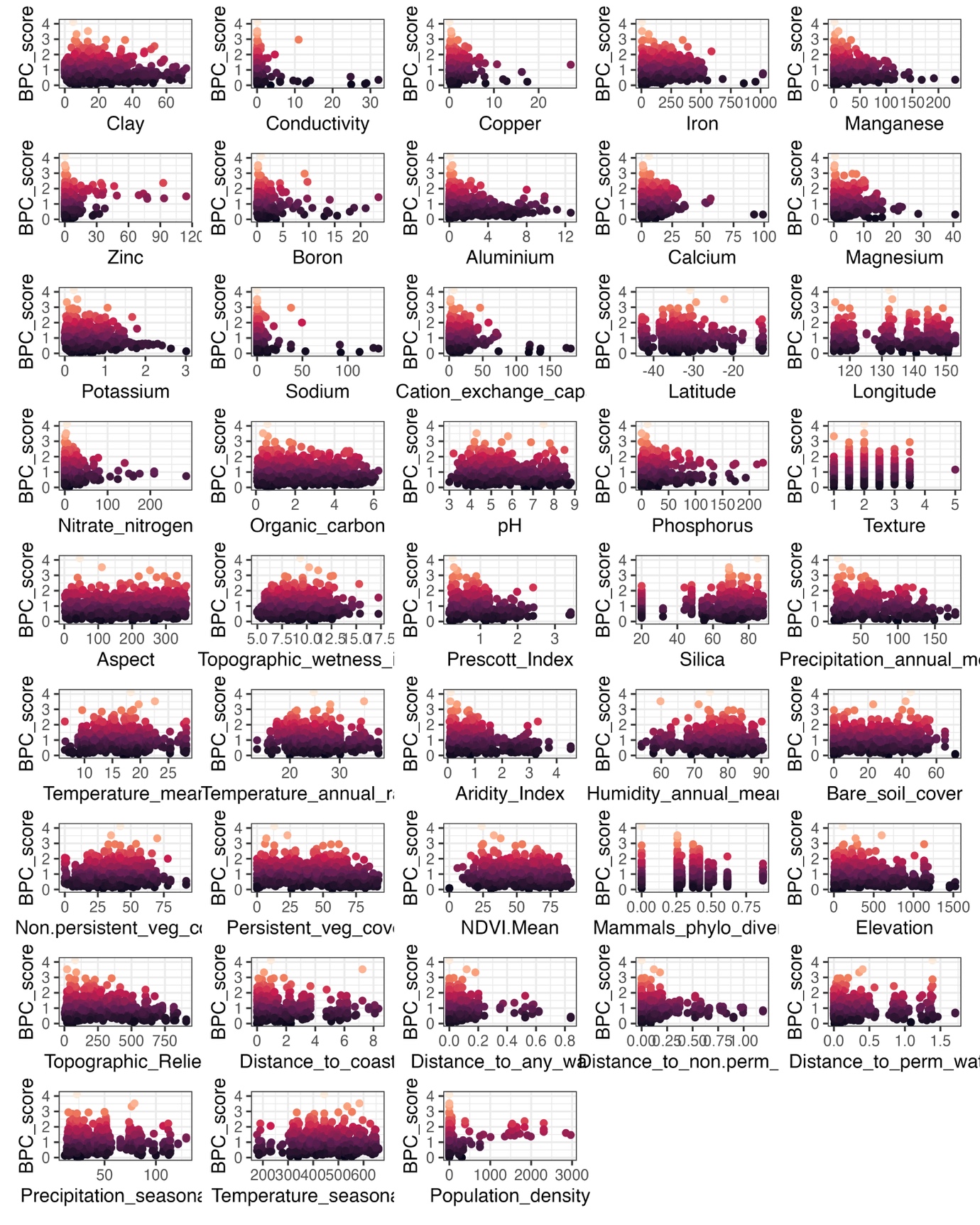

**Supplementary Figure 3. Scatterplots of ecological covariates with BPC_16S_ scores.**

Scatterplots of BPC_16S_ scores against 43 continuous ecological predictor variables.

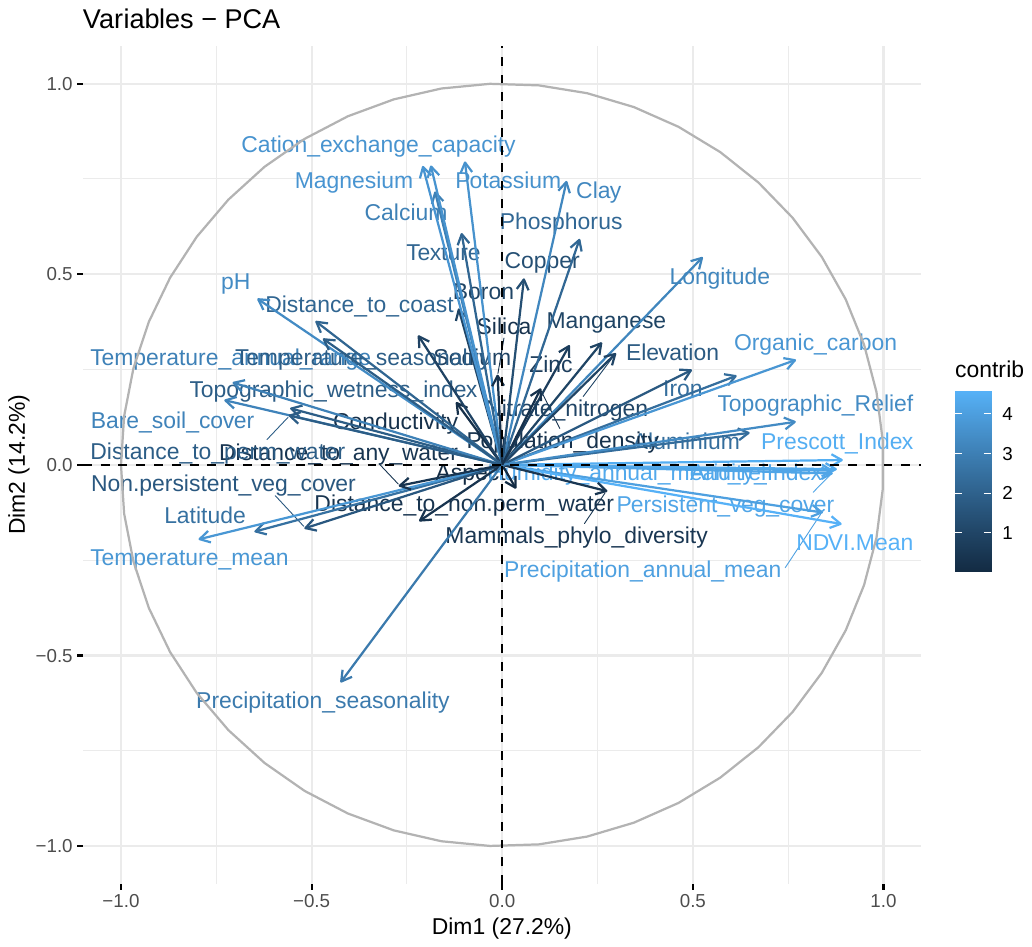

**Supplementary Figure 4. PCA variables plot from analysis of Australian 16S rRNA soil samples.**

Contributions of 43 continuous predictor variables to the top two principal components from principal component analysis. Longer arrows indicate greater contribution. “Dim1” generally represents climatic influences, and “Dim2” generally represents soil fertility and cation content.

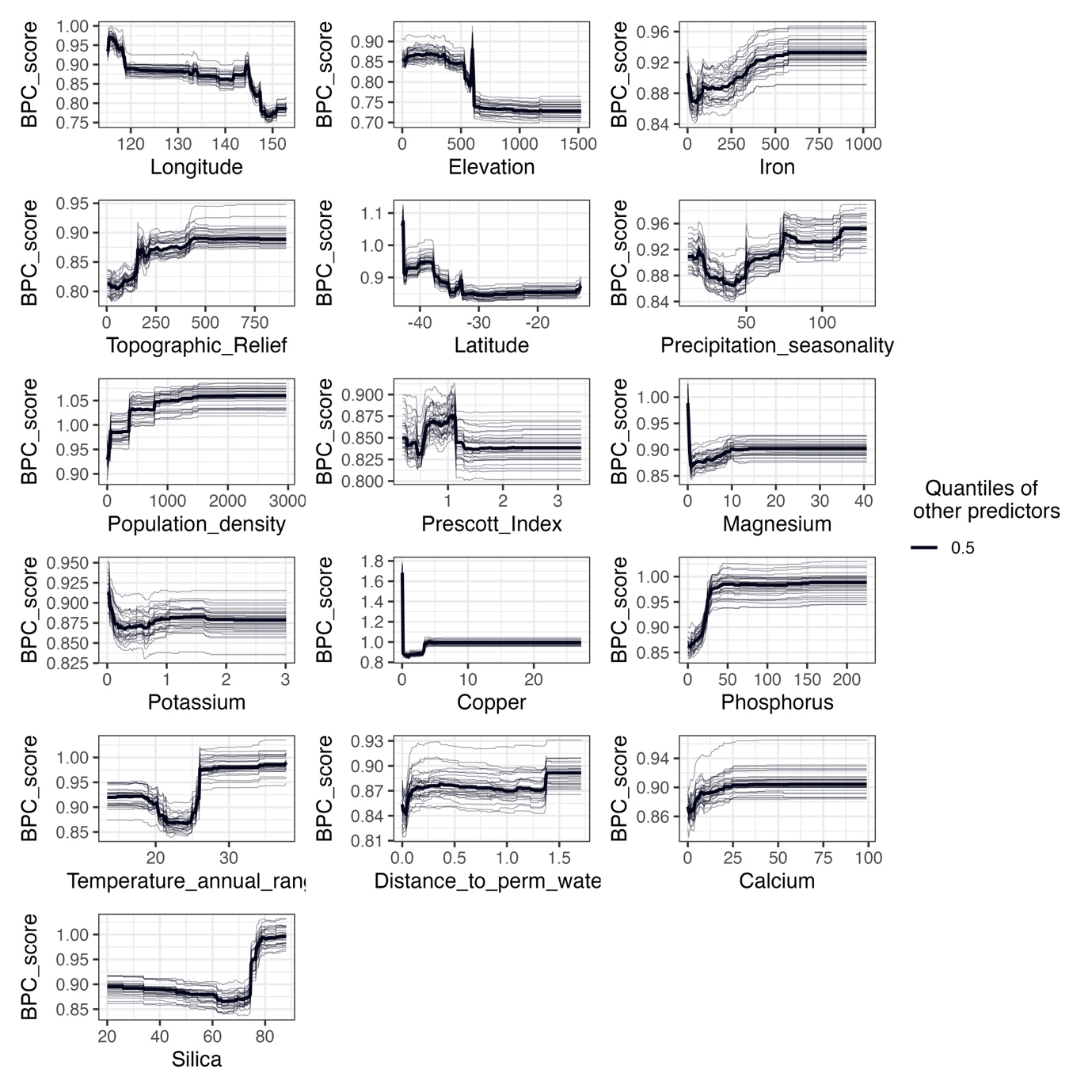

**Supplementary Figure 5. Partial dependence plots from Random Forest modelling with Australian 16S rRNA soil samples.**

Partial dependence plots from Random Forest modelling with Australian 16S rRNA soil samples and ecological covariates. Only covariates with a variable importance score above the median are shown. Trendlines indicate changes in BPC_16S_ score response while maintaining centred values of ecological covariates.

**Supplementary Figure 6. Anthrome-based world map of downloaded soil and land-based sediment samples.**

Visualization of variation in BPC_meta_ scores of global soil and land-based sediment samples using an anthrome-based world map. *n*=2850. BPC_meta_ scores indicated by point colors. Anthrome classification legend shows anthrome types (“Intensive”, “Cultured”, and “Wild”), levels (“Dense Settlements”, “Villages”, “Croplands”, “Rangelands”, “Cultured”, and “Wildlands”), and classes (e.g., “Urban” and “Residential woodlands”).

Supplementary Tables

The following datasets generated during and/or analysed in the current study are available on figshare at <https://figshare.com/s/3ae2f1327a11f91793d8>.

**Supplementary Table 1. Descriptions of genes considered for analyses of global metagenomic samples.**

This table lists and describes the genes within the butyrate synthesis pathway considered for analysis of global metagenomic samples. The table includes the EC number of the associated enzymes, the general function of each enzyme within the butyrate synthesis pathway, and reason for inclusion or exclusion in our analyses.

**Supplementary Table 2. Gene counts of *buk* and *atoA* genes among metagenomic samples.**

This table shows *buk* counts and *atoA* counts on two separate sheets for each bacterial genome from the IMG/M database with at least one of either gene. Mean gene count of *buk* was 1.180044. Mean gene count of *atoA* was 2.113735. “Status” column shows genome completion: “F”=Finished, “P”=Permanent draft, “D”=Draft.

**Supplementary Table 3. Downloaded metagenomic data and metadata from 22,593 samples.**

This table provides the source download data from 22,593 metagenomic samples. The first sheet is the combined data, and subsequent sheets show separately the gene count data for *atoA*, *buk*, and *pheS* and sample metadata.

**Supplementary Table 4. Downloaded metagenomic sample data distributed into six general source categories.**

This table shows the distribution of metagenomic sample data into six general source categories. Each sheet in the workbook has a separate category: Human, Non-human animal gut, Plant, Soil, Aquatic, and Agro-Industrial. Columns (A)-(K) are gene data and calculations of BPC_meta_ scores. Columns (L)-(BD) are sample metadata.

**Supplementary Table 5. Set of 118 putative butyrate-producing bacteria species and their taxonomic data.**

This table lists the set of 118 putative butyrate-producing bacterial species (Column F) and their taxonomic families (Column G), with the proportion of species within each family that are putative butyrate producers (Columns A-D). The species set was developed from Vital et al^31^, with taxonomic and proportion data derived from Genome Taxonomy Database (GTDB).

**Supplementary Table 6. Ecological covariates used in the analyses of Australian 16S rRNA soil samples.**

This table describes each of 49 ecological SCORPAN predictor variables. The predictor variables include 43 continuous variables and 6 categorical variables. For each variable, the table lists the download source, ecological range/units, type of data, download date, and general description.

**Supplementary Table 7. Gene data, metadata, and covariate data for 1,331 Australian 16S rRNA soil samples.**

This table provides the data used in analysis of 1,331 Australian 16S rRNA soil samples with 49 ecological SCORPAN covariates. Samples with incomplete data were removed.

**Supplementary Table 8. Data for each of five land type clusters generated from Australian 16S rRNA soil sample analysis.**

This table provides data for each land type cluster derived from *k*-means clustering. Data for the 43 numeric predictor variables and sample BPC_16S_ scores are included. All listed data are medians.

**Supplementary Table 9. Data for categorical variables generated from Australian 16S rRNA soil sample analysis.**

This table provides BPC_16S_ data for each category of the categorical predictor variables: land cover, land use, anthropogenic biome, vegetation types, and major vegetation subgroups. The table includes the count, mean BPC_16S_ score, median BPC_16S_ score, and standard deviation and is sorted by descending medians.
